# Single neuron firing cascades underlie global spontaneous brain events

**DOI:** 10.1101/2021.01.22.427798

**Authors:** Xiao Liu, David A. Leopold, Yifan Yang

## Abstract

The resting brain consumes enormous energy and shows highly organized spontaneous activity. To investigate how this activity is manifest among single neurons, we analyzed spiking discharges of ∼10,000 isolated cells recorded from multiple cortical and subcortical regions of the mouse brain during immobile rest. We found that firing of a significant proportion (∼70%) of neurons conformed to a ubiquitous, temporally sequenced cascade of spiking that was synchronized with global events and elapsed over timescales of 5-10 seconds. Across the brain, two intermixed populations of neurons supported orthogonal cascades. The relative phases of these cascades determined, at each moment, the response magnitude evoked by an external visual stimulus. Furthermore, the spiking of individual neurons embedded in these cascades was time locked to physiological indicators of arousal, including local field potential (LFP) power, pupil diameter, and hippocampal ripples. These findings demonstrate that the large-scale coordination of low-frequency spontaneous activity, which is commonly observed in brain imaging and linked to arousal, sensory processing, and memory, is underpinned by sequential, large-scale temporal cascades of neuronal spiking across the brain.

## Introduction

The brain at rest exhibits slow (<0.1 Hz) but highly organized spontaneous activity as measured by functional magnetic resonance imaging (fMRI)^1,2^. Much of research in this area has utilized the temporal coordination of these signals to assess the functional organization a large number of brain networks. In recent years, however, new attention has been directed to a less studied aspect of this signal, namely the conspicuous and discrete spontaneous events that occur simultaneously across the brain^3–5^. These global resting-state fMRI events appear to reflect transient arousal modulations at a timescale of ∼10 seconds^4,6^ and also to be closely related to activity among clusters of cholinergic projection neurons in the basal forebrain^4,5^.

The nature of global brain events is of great interest, as is their spatiotemporal dynamics. Some evidence suggests they take the form of traveling waves, propagating coherently according to the principles of the cortical hierarchy^7–9^, and shaping functional connectivity measures important for assessing the healthy and diseased brain^9–11^. Other work has linked such global activity to phenomena as varied as modulation of the autonomic nervous system^12–15^, cleansing circulation of cerebrospinal fluid in the glymphatic system^16–19^, and memory consolidation mediated by hippocampal sharp-wave ripples^20,21^. In general, the global activity measured through brain imaging appears coordinated over timescales of seconds with a range of other neural and physiological events^12,13,15,21–23^. In a few cases, the relationship between local and global neural events has been studied using simultaneous measurements. For example, brain-wide fMRI fluctuations and local field potential (LFP) power changes are locked to the issuance of hippocampal ripples^21,24^. However, very little is understood about the extent to which single neurons participate in the expression and coordination of global spontaneous events of the seconds timescale. To approach this topic, recent technological advances have made it possible to track and compare the spiking activity of a large number of isolated neurons recorded simultaneously across multiple brain areas.

A few recent studies utilizing high-density neuronal recording^25^ have accumulated initial evidence suggesting a close relationship between the brain state and neuronal population dynamics^26–28^. A large proportion of neurons, regardless of their location, showed strong modulations in their discharging rate that were coordinated in time with physiological arousal measures^28^, across thirsty and sated states^26^, and during exploratory and non-exploratory behaviors^27^. Nevertheless, these studies leave open the question of how neuronal population dynamics are organized at a finer timescale of seconds surrounding spontaneous global events during immobile rest, and whether and how such dynamics are coincident with arousal modulations, hippocampal ripples, and sensory excitability. To investigate this topic, we examine the spiking activity recorded from large neuronal populations of neurons in immobilized mice, focusing on their seconds-scale coordination with global events and with one another. We further studied the impact of this spontaneous spiking on the magnitude of visually evoked responses, and its time locking with other physiological signals related to arousal, such as LFP changes, hippocampal ripples, and changes in pupil diameter.

## Results

We analyzed neural activity, pupillary changes, and locomotor activity obtained from the Visual Coding – Neuropixels project of the Allen Brain Institute^29^. We focused on the spontaneous spiking activity of ∼10,000 neurons from 44 brain regions of 14 mice (730 *±* 178 neurons per mouse, mean*±*SD) sustained during periods of immobile rest (**Fig. 1A** and **Table S1**).

**Figure 1.**
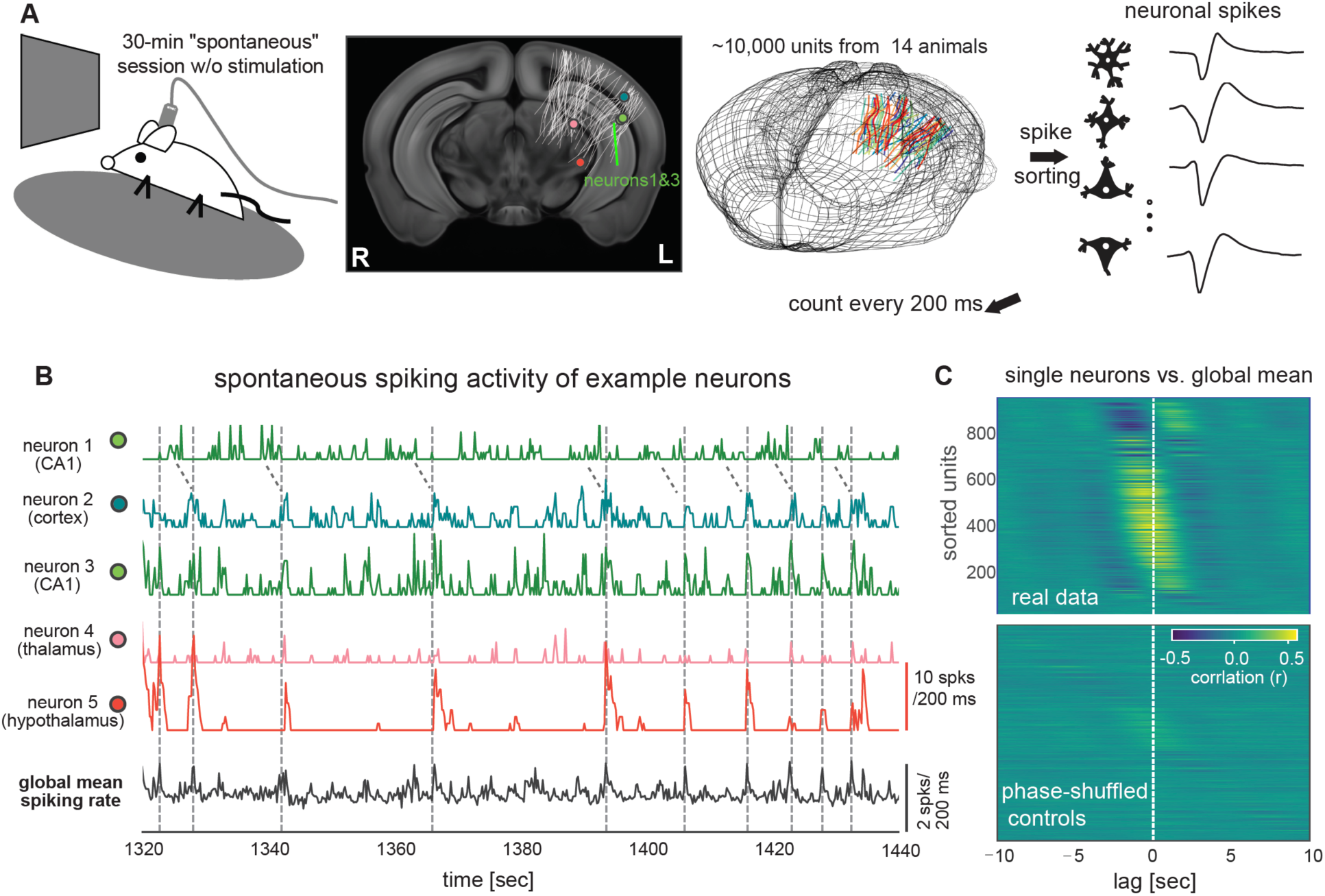
Neuronal population is engaged in global activity of the seconds timescale during immobile rest. (A) Illustration of the experimental setup of the Visual Coding – Neuropixels project that included a 30-min “spontaneous” session without any stimulation (top-left). The locations of 6171 channels on 79 probes from 14 mice were collapsed along the anterior-posterior direction and mapped onto a middle slice of a mouse brain template and also shown in a 3D representation of mouse brain (the second and third columns). (B) Spontaneous spiking rate of five example neurons from the hippocampus (CA1), visual cortex (VISal), thalamus (LP), and hypothalamus (ZI) of a representative mouse. Their locations were marked in (A). The neurons 1 and 3 were recorded by two channels with a distance of 14um. The bottom trace is the global mean spiking rate of all 930 surveyed neurons. (C) Cross-correlation functions between individual neurons’ spiking rate and the global mean spiking rate (reference) during the stationary period from the representative mouse (top). The same cross-correlation functions were computed after phase-shuffling the real data (bottom).

### Global coordination of spontaneous spiking activity during rest

The 30-min “spontaneous” sessions were characterized by two distinct behavioral states, with mice either running or remaining still. The two states were characterized by distinct patterns of neural and behavioral signals (**Fig. S1A**), with the immobile rest associated with reduced pupil diameter (*p* = 2.2*×*10^−7^, *N* = 14; paired t-test), pupil motion (*p* = 8.2*×*10^−7^), eye-area height (*p* = 0.0008) (**Fig. S1B**), and mean spiking rate (*p* = 1.1*×*10^−5^), all of which suggested a less aroused state.

Close inspection of neural activity during immobile rest revealed bouts of elevated spiking in majority of neurons across distant brain areas that included the visual cortex, hippocampus, thalamus, and hypothalamus (**Fig. 1B**). These ubiquitous and coarsely synchronous events occurred aperiodically, with typical intervals in the range of 5-20 seconds, although the spiking bursts observed between a given pair of neurons were frequently shifted in time (see neurons 1 and 3 in **Fig. 1B**). These quasi-synchronized firing events led to large peaks in the global mean spiking rate (here and throughout, “global” refers to the surveyed population) of all survey neurons (dashed lines, **Fig. 1B**). As a result, the stationary period was characterized by a large fluctuation in the global mean spiking rate and a high level of global synchronization as measured by the Fano factor (**Fig. S1C**)^30,31^.

To understand the temporal relationship between the firing of individual neurons and the average population dynamics during immobile rest, we correlated each neuron’s spiking activity with the global mean spiking rate. Cross-correlation functions were computed from lowpass filtered (< 0.5 Hz) spiking rate time courses in order to focus on slow modulations. This analysis revealed two main findings. First, the spontaneous firing of the majority of surveyed neurons (70.4% *±* 8.5%, across the 14 mice; **Fig. S1D**), regardless of the location (**Fig. S2**), were significantly (*p* < 0.05; permutation test as compared to spectral-profile-preserved but phase-shuffled controls) correlated with the global mean spiking rate (**Fig. 1C**). Second, the peak correlations for individual neurons spanned time lags ranging between *±* 5 seconds (**Fig. 1C** and **S1E**), demonstrating a unique and consistent time delay between a single neuron’s firing and the global spiking activity.

### Sequenced activity cascades underlie spontaneous resting activity

Given that individual neurons appeared to be entrained into the global activity with distinct time delays, we thus examined if the structured patterns of sequential activations, which have been observed in resting-state fMRI signals^7,9^, exist in populational spiking data. We adapted a delay-profile decomposition method, which has been validated with simulation and successfully extracted propagating patterns of resting-state fMRI signals^7^, to achieve this goal. Briefly, the spiking data was cut into time segments based on troughs of the global mean signal, and a delay profile was computed for each segment to describe relative delays of each neuron’s spiking activity based on the centroid of spiking time. A singular value decomposition (SVD) was then applied to all delay profiles to extract their first principal component (i.e., the principal delay profile) that would represent the major direction of sequential activations if any exists (**Fig. 2A**; see Methods for details). Applying this method to the spiking data extracted a principal delay profile that accounts for a significant portion of delay profile variance (**Fig. S3**).

**Figure 2.**
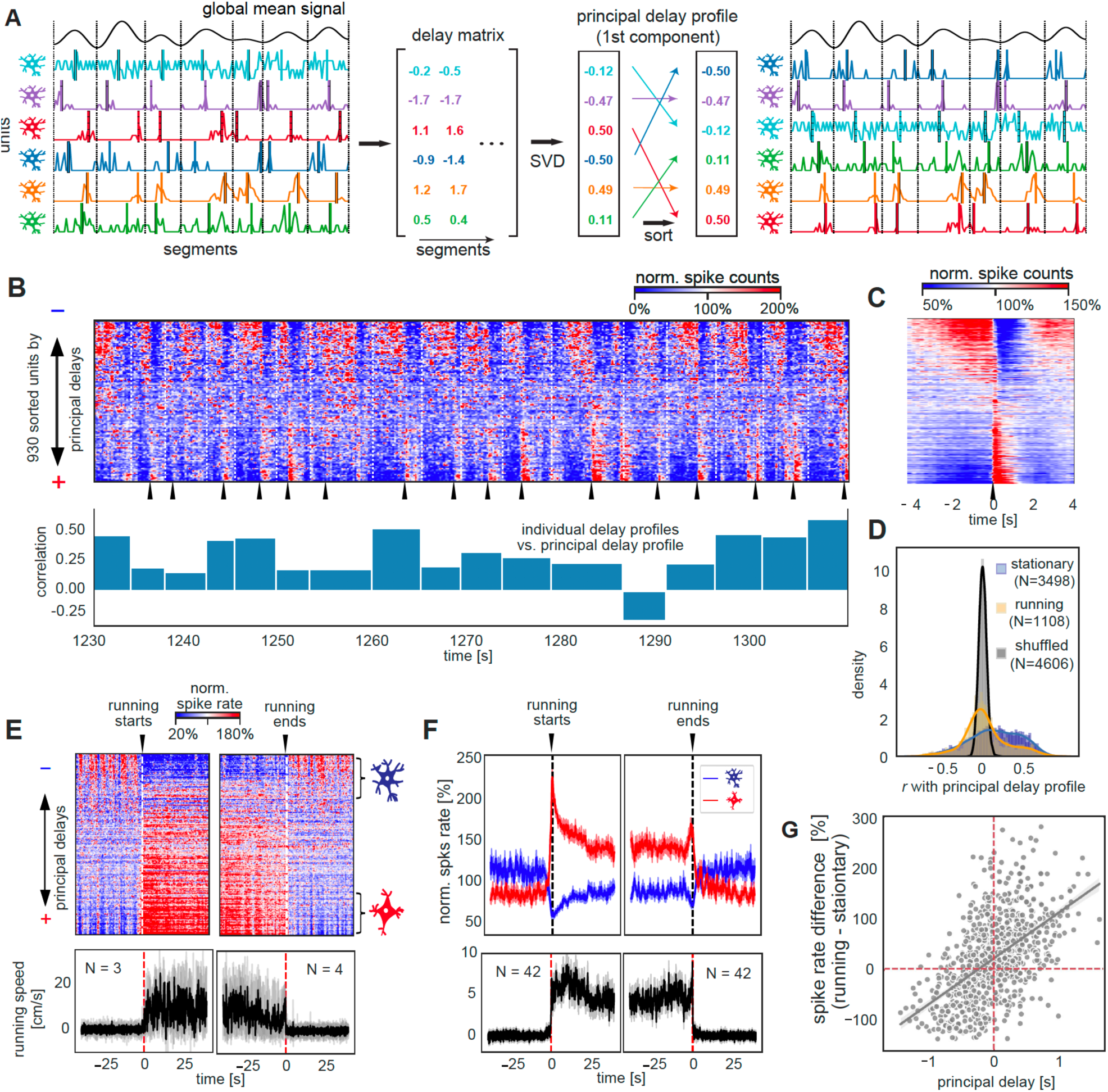
Sequential activations between two distinct neuronal ensembles define the resting-state neuronal dynamics. (A) Illustration of the delay-profile decomposition method to identify and extract sequential activation pattern in neuronal spiking data. The spiking data were divided into segments according to troughs of the filtered (<0.5 Hz) global mean signal. A delay profile was computed for each segment based on the centroid of spiking activity and then assembled to a delay matrix. The singular value decomposition (SVD) was applied to the delay matrix to extract the principal delay profile (i.e., the first principal component) that represents the major direction of sequential activations across neurons. Sorting neurons according to this principal delay profile allows us to visualize sequential activations of neurons. (B) An 80-sec example of spontaneous spiking activity of the representative mouse with all 930 neurons sorted according to the principal delay profile (top). The sharp activations of the positive-delay neurons were marked by black arrows. The bar plot (bottom) shows the correlations between the principal delay profile and the delay profile of individual time segments, whose boundaries were marked by white dashed lines. (C) The mean pattern of sequential activation of the representative mouse was obtained by aligning and averaging the time segments of sequential activation according to the sharp activations of the positive-delay neurons. (D) Distributions of the correlations of individual delay profiles with the principal delay profile. They were summarized for the stationary (blue) and running (orange) periods, as well as the randomly shuffled (black) individual delay profiles. (E) Spiking activity (top) and running speed (bottom) at the transitions of running and stationary states for the representative mouse. The black and gray lines represent the mean and individual traces respectively. (F) Spiking activity of the negative- and positive-delay neurons (top) and running speed (bottom) at the transitions of running and stationary states summarized across all 14 mice. The thick lines represent the mean whereas shadow regions denote areas within one S.E.M. (G) The principal delay value of individual neurons of the representative mouse is significantly correlated (*r* = 0.54, *p* = 2.5*×*10^−71^) with their spiking rate change across the running and stationary states.

Sorting the neuronal spiking time courses according to the derived principal delay profile revealed sequenced spiking cascades, with individual neurons exhibiting a gradation in their temporal shifts relative to shared global events (**Fig. 2B**, top). Each cascade was repeated with each cycle of the global mean signal, with individual neurons having fixed temporal lags. These cascades were most prominent during periods of immobile rest. Across time, 51.1%*±*13.7% of stationary time segments exhibited a significant (p<0.001, permutation test) positive correlation with the principal delay profile. The running time segments showed much lower correlations (*p* = 2.6*×*10^−35^, Mann-Whitney U test) with the principal delay profile (**Fig. 2D**), with relatively strong correlations occurring mostly at short epochs of immobility between running, which were classified to the running state (see Methods for details).

To better visualize the average temporal structure of each neuron in the population, we aligned to sharp rate increases evident in positive-delay neurons (black arrows in **Fig. 2B**, see Methods for the definition of the positive- and negative-delay neurons). This analysis revealed the sequence of events in the neural population (**Fig. 2C**), beginning with the increased spiking of some negative-delay neurons, with gradual recruitment of additional neurons, followed by an abrupt transition in which the negative-delay cells ceased firing and the positive-delay cells showed a sudden firing increase, which decayed within ∼1s. This pattern was highly reproducible across animals (**Fig. S4**).

Analysis of the same neurons during periods of active running revealed that the negative- and positive-delay groups constituted different functional subpopulations. Specifically, the spiking rate of negative-delay neurons was suppressed during running, whereas that of positive-delay neurons was substantially increased (**Fig. 2E**-**2F**). Across the population, correlation between a neuron’s principal delay value at rest and its spike-rate changes during running was robust (*r* = 0.55 *±* 0.08, *p* = 2.9*×*10^−12^), with negative delays largely associated with spiking reduction during running (**Fig. 2G**).

To summarize, activity during rest was composed of highly structured, repeating patterns of sequenced spiking activation that fell into two distinct neuronal groups with ostensibly different functional roles during behavior.

### Spatial organization of the negative- and positive-delay neurons

Negative-delay and positive-delay neurons were often spatially intermingled within the same brain areas, suggesting that the expression of spontaneous events at the single neuron level was not that of a coherent spatiotemporal wave (**Fig. 3A**). While cortical areas had a roughly even balance of leading vs. lagging neuronal activity, CA1 and dentate gyrus (DG) of the hippocampus were dominated by negative delays (*p* < 2.8*×*10^−29^, Bonferroni corrected) whereas the thalamus was dominated by positive delays (*p* < 3.0*×*10^−12^, Bonferroni corrected) (**Fig. 3B**, left). These systematic biases stemmed from unbalanced proportions of negative-delay and positive-delay neurons: the CA1 and DG regions contain significantly more (*p* < 6.5*×*10^−29^, proportion z-test, Bonferroni corrected) negative-delay neurons whereas the thalamic structures have significant less (*p* < 9.8*×*10^−53^), as compared with the overall distribution (**Fig. 3B**, right).

**Figure 3.**
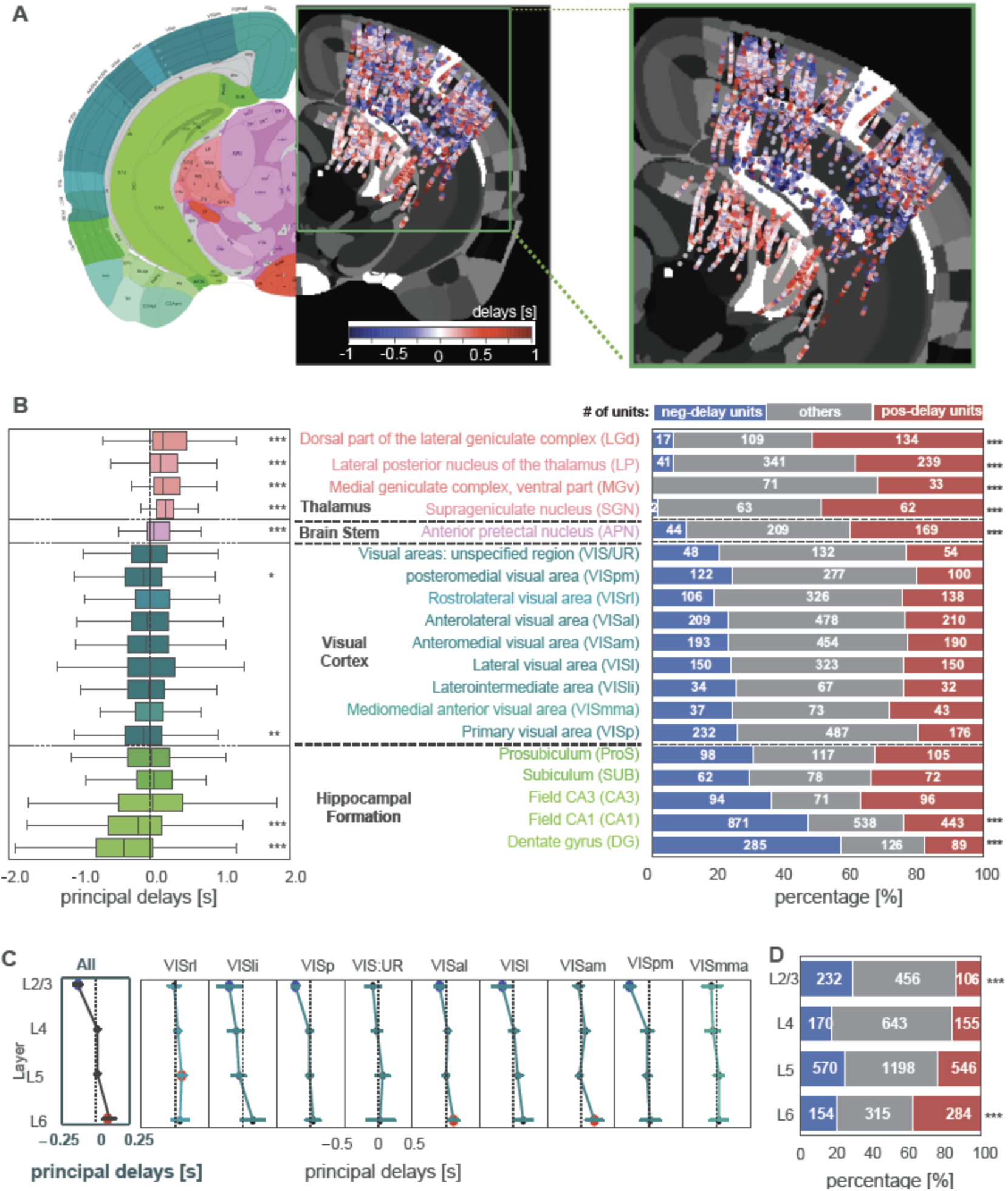
Spatial organization of the negative- and positive-delay neurons. (A) A map of the principal delay values. The principal delay values were averaged across units detected by the same channel, combined across all 14 animals, collapsed along the anterior-posterior direction, presented on an axial brain slice in the middle of recording sites, and shown along with a corresponding mouse brain atlas (left; image credit: Allen Institute). The recording areas were amplified and shown on the right. (B) A boxplot showing the region-specific distributions of the principal delay values (left) is displayed together with a bar plot showing the percentages of three groups of neurons with distinct delays (right). (C) The distributions of the principal delays across different cortical layers at various visual regions (pooled results in the first column). Error bars indicate 95% conference intervals of the mean delay. Brain regions with a mean delay significantly (*p* < 0.05) different from zero are marked by colored circles (blue: negative; red: positive). (D) The percentages of the three types of neurons at different cortical layers. Asterisks represent the significance level (Bonferroni corrected: ∗, 0.01< *p* < 0.05; ∗∗, 0.001< *p* < 0.01; ∗∗∗, *p* < 0.001.

Similar biased distributions of the two types of neurons were also observed across cortical layers (**Fig. 3A**). Despite an overall balanced distribution of the principal delay values in the cortex, the mean delay value of layers 2/3 (L2/3) neurons was significantly (*p* = 7.5*×*10^−17^) lower than zero whereas that of layer 6 (L6) neurons was significantly positive (*p* = 1.2*×*10^−5^) (**Fig. 3C**). Accordingly, relative proportions of the leading and lagging neurons was significant different in the L2/3 (*p* = 6.1*×*10^−12^) and L6 (*p* = 2.9*×*10^−11^) layers as compared with the overall distribution (**Fig. 3D**). We further examined the sub-second time delay among the leading/lagging neurons of different visual cortical layers with those in the hippocampus and thalamus. According to the peak correlation offset, the CA1 and DG leading neurons preceded the visual L2/3 leading neurons by 138 *±* 126 ms (*p* = 0.0019) in their spontaneous firing, whereas the visual L6 lagging neurons led the thalamic lagging neurons by 169 *±* 160 ms (*p* = 0.0024) (**Fig. S5**).

Together, these findings suggest that the observed spontaneous activation sequences are expressed earliest in the hippocampus, and then propagate to the cortex, first affecting the supragranular layers, followed by the infragranular layers and then the thalamic projection targets.

### Sequential activation, arousal, and hippocampal ripples

We further examined changes of arousal-related measures across the cycle of global sequential activation event, since the global resting-state fMRI activity has been linked to transient arousal modulations^4^. First, we found that pupil diameter fluctuations were phase locked to the sharp activation of lagging positive-delay neurons (**Fig. 4A**, the second row). Second, we found that the cortical delta-band (0.5-4 Hz) activity demonstrated a concurrent decrease in its power, resulting slow modulations of seconds timescale (**Fig. 4A**, the third row). For both of these arousal indicators, the average patterns within (**Fig. 4B**) and across animals (**Fig. 4C**) showed strong and distinct modulations across the cycle of sequential activation. The delta-power changes closely followed the temporal dynamics of the negative-delay neurons with a mean delay of 339 *±* 199 ms (*p* = 2.5*×*10^−5^), as estimated by the peak correlation offset (**Fig. 4D**, left), whereas the pupil diameter changes followed the spontaneous firing of the positive-delay neurons by 1046 *±* 118 ms (*p* = 6.2*×*10^−15^) (**Fig. 4D**, right). These changes were spontaneous, with the animal at rest, and thus could not be explained by potential residual motion of animals (**Fig. 4A-C**, the bottom row).

**Figure 4.**
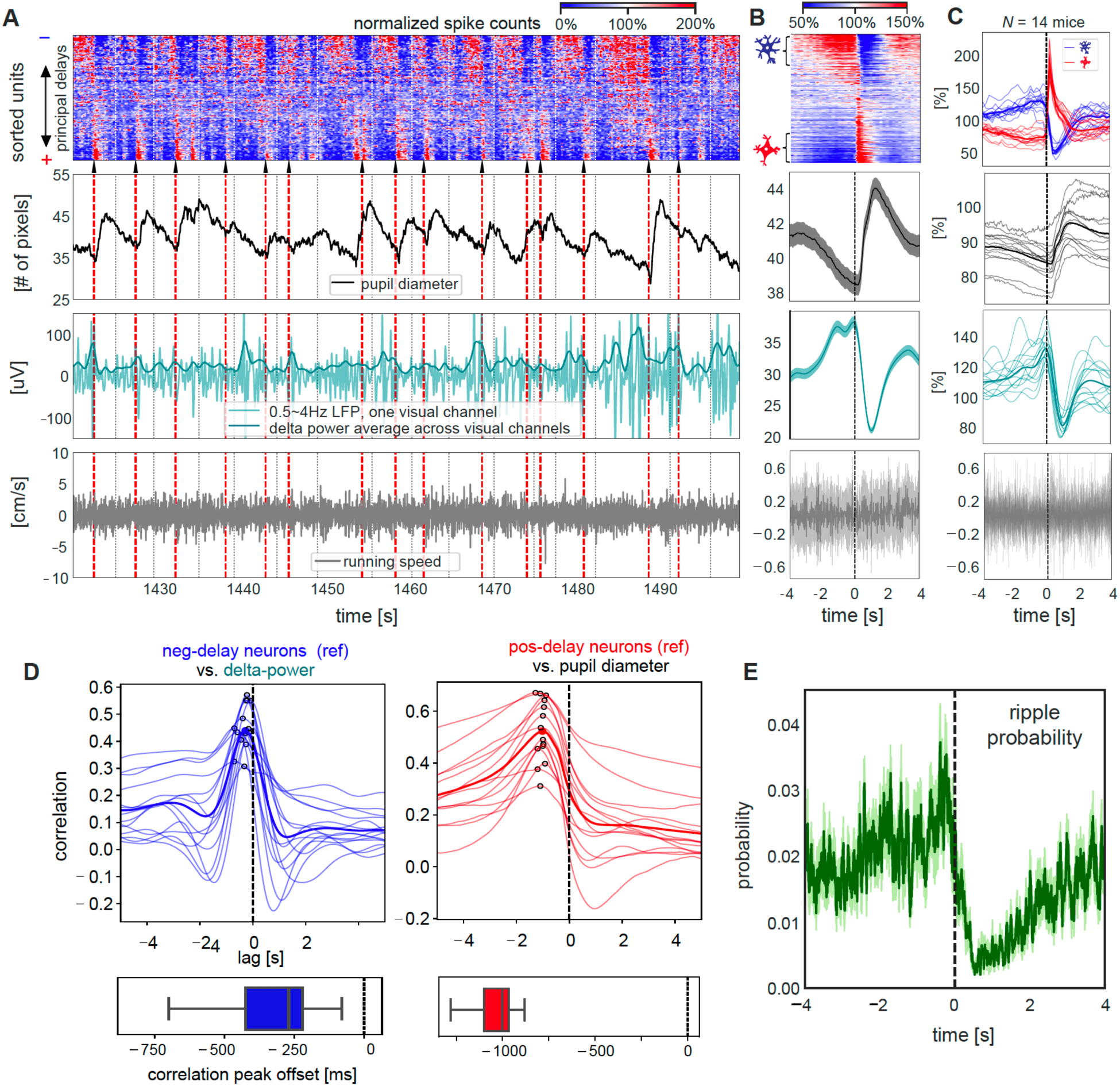
Sequential activation pattern is accompanied by seconds-scale modulations of arousal measures and hippocampal ripples activity. (A) An 80-sec section of various neuronal and behavioral signals from the representative mouse. Sequential activation patterns of the spiking activity (top) are associated with changes in pupil diameter (the second row) and delta (0.5-4 Hz) activity (the third row), but not running speed (the bottom row). White/black dashed lines denote time segment boundaries. Black arrows and red dashed lines mark the sharp activation of the positive-delay neurons, according to which the time segments were aligned and averaged. The time segments with delay profile positively correlated with the principal delay profile were aligned and averaged according to the sharp activation of the positive-delay neurons within the representative mouse (B) and across all 14 mice (C). For the group results (C), the spiking dynamics of the negative- and positive-delay neurons were shown, and the thicker, darker lines represent the mean curves whereas the thinner, lighter lines denote the results from individual animals. For the results of the representative mouse (B), the thicker, darker lines represent the mean whereas the shadowed regions denote the regions within one SEM across the 14 animals. (D) Cross-correlation functions were computed between the spiking activity of negative-delay neurons (reference) and the cortical delta-power (left), as well as between the positive-delay neurons’ spiking dynamics (reference) and the pupil diameter (right), during the stationary period. Thin and thick lines present the results of individual animals and their average respectively, and the peak correlations are marked by hollow circles. The boxplots (bottom) show the distribution of the time offset of peak cross-correlations. (E) The hippocampal ripple events were aligned and averaged across the time segment of sequential activations during the stationary period, similar to the other neural and behavioral signals in (C). The shadow regions represent areas within one SEM across the 14 animals.

We also examined how this global sequential activation might be related to hippocampal ripples, which have been shown to co-modulate with cortical delta-power^24^, pupil microdilations^32^, and brain-wide fMRI changes^21^ on a similar seconds timescale. Adapting an existing algorithm^33^ (see Methods for details), we detected 606 *±* 184 hippocampal ripple events (average life: 46.2 *±* 17.8 ms) per mouse/session from LFPs recordings in the CA1 region, and they were predominantly during the stationary period as expected (**Fig. S6**). The occurrence probability of these ripple events was strongly modulated across the cycle of sequential activation (**Fig. 4E**). Specifically, it closely followed spiking dynamics of the negative-delay neurons and occurred with a probability close to zero during the sharp activation of the lagging positive-delay neurons. Altogether, the sequential activation of two distinct neuronal populations were orchestrated with the seconds-scale modulation of both arousal measures and hippocampal ripple activity.

### Ongoing activity of the two neuronal ensembles affects stimulus-evoked responses

Given the prominent role of the negative- and positive-delay neurons in resting-state brain dynamics, we investigated whether their activity state might affect responses to external sensory stimuli^34–36^. To achieve this goal, we used spiking data in another 30-min session with drifting grating stimulation (**Fig. 5A**). We quantified the pre-stimulus state (−0.2-0 sec to the stimulus onset) by the firing rate difference between the negative-delay and positive-delay neurons (those responding to the drifting grating were excluded). The response amplitude of the third group of neurons, which were significantly activated by the drifting gratings (those with significant negative or positive principal delays were excluded), was quantified for both an early onset phase (0-0.4 sec) and a later steady-state period (0.4-2 sec) during stimulation (**Fig. 5B**).

**Figure 5.**
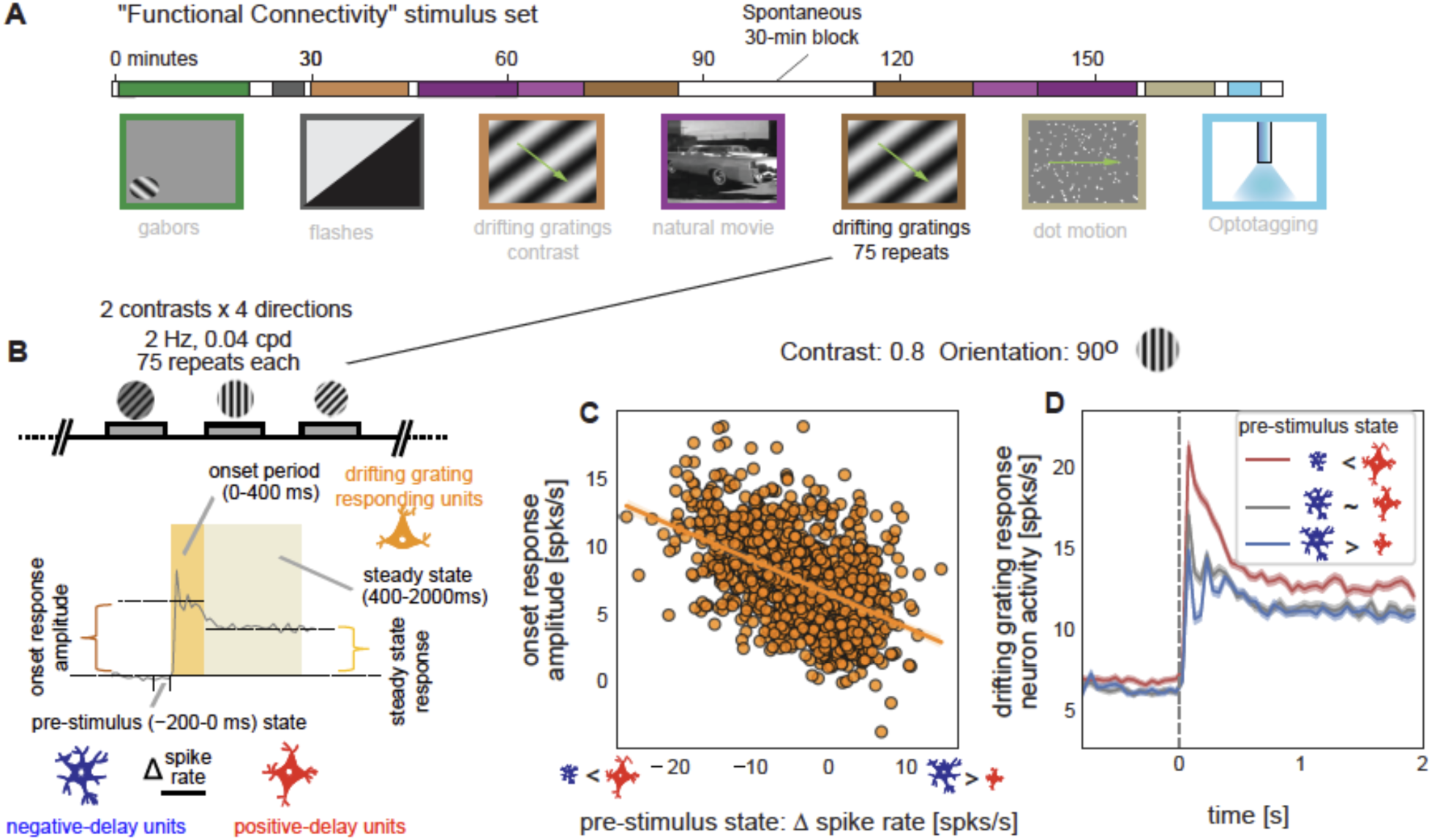
Ongoing activity of the two neuronal ensembles predicts the amplitude of visual responses. (A) A diagram showing the “Functional Connectivity” stimulus set of the “Visual Coding Neuropixels” project, which included a 30-min session with drifting gratings stimuli. (B) Illustration of the definition of the pre-stimulus state and the response amplitude to drifting gratings stimuli (image credit: Allen Institute). The pre-stimulus state was defined as the spiking rate difference between the negative- and positive-delay neurons right before the stimulus onset (−200-0 ms), whereas the response amplitude was defined as the spiking rate change of the drifting-grating responding neurons during the onset (0-400 ms) and steady-state (400-2000 ms) stimulation periods as compared with the baseline period (−800-0 ms) without stimulation. (C) The pre-stimulus state of the negative- and positive-delay neurons is significantly correlated with the onset response amplitude of the drifting grating responding neurons to a drifting gratings stimulus (orientation: 90 degree; contrast: 0.8) across 1050 trials pooled from all 14 mice (*r* = −0.50, *p* = 3.9*×*10^−66^) (D) Spiking rate of the drifting gratings responding neurons were averaged within three groups of trials divided based on the pre-stimulus state: the bottom 350 trials (red) with the positive-delay neurons more active, the middle 350 trials (gray) with similar spiking rate between the negative- and positive-delay neurons, and the top 350 trials (blue) showing more active negative-delay neurons. Shadows represent regions within *±* one SEM across trials.

As shown by the example for a specific stimulus (90° and 0.8 contrast), the onset response amplitude of the drifting-grating responding neurons to the visual stimulus was strongly dependent on the pre-stimulus state of the negative- and positive-delay neurons (*r* = −0.50, *p* = 3.9*×*10^−66^) (**Fig. 5C**). The neurons’ response to identical drifting grating patterns was often several fold higher when positive-delay neurons were more active right before simulation (**Fig. 5C**). The association was reproducible for all eight stimulus types (*r* = −0.52 *±* 0.03, *p* = 3.9*×*10^−10^) and became weaker (*p* = 8.7*×*10^−5^) but still significant (*r* = −0.32 *±* 0.05, *p* = 5.6*×*10^−7^) for the steady-state response (**Fig. S7**). Consistent with these correlative analyses, the drifting-grating responding neurons showed distinct dynamics, particularly during the onset phase, in trials with distinct pre-stimulus states (**Fig. 5D**). Overall, the pre-stimulus activity of the negative- and positive-delay neurons accounted for a significant portion of trial-to-trial variability in neuronal response to identical visual stimuli.

## Discussion

Here we showed, using large-scale single-unit recordings in mice, that a surprisingly large proportion (∼70%) of surveyed neuronal population, regardless of their location, were entrained to slow (∼5-10 seconds) but highly structured spontaneous brain activity at immobile rest. Activity relative to global events was characterized by sequential activations between two distinct neuronal ensembles that showed orthogonal modulations across running/stationary states. The cortical delta-band power and pupil diameter showed similar, but slightly delayed, dynamics following spontaneous firing of the leading, rest-active neurons and the lagging, running-active neurons respectively, leading to their strong modulations across the sequential activation cycle. In addition, the occurrence of the hippocampal ripples was tightly linked to this global sequential activation and closely followed the dynamics of leading rest-active neurons. Lastly, the pre-stimulus activity of the two neuronal ensembles accounts for the trial-to-trial variability in neuronal responses to subsequent visual stimuli. Overall, these findings provide a novel, single-cell perspective on resting-state seconds-scale brain dynamics at the cellular level and suggest a specific entity that may link arousal and memory functions of the brain, which are often concurrently impaired in neurodegenerative diseases.

While our recordings were extensive, they only covered a subregion of the brain, thus for cortical and subcortical structures that we did not analyze, we cannot know for certain whether the same cascades are present. However, there is some reason to believe that these signals are, to a first approximation, “global”. First, all 44 surveyed brain regions from various cortical and subcortical structures, e.g., the visual cortex, thalamus, brainstem, and hippocampus, in our study were replete with neurons that were active during these widespread events (**Fig. S2**). Second, the recent studies reporting ensembles of neurons showing opposite modulation across distinct brain states^26–28^, including the running and stationary state^28^, were similarly found at all recording sites across an even wider span of brain regions. Those neuronal ensembles are likely overlapped with the negative- and positive-delay neurons found during rest in the present study. Third, this global neuronal spiking activity shares important features with the global resting-state fMRI signal, which has been shown to involve brain-wide changes^4^. For example, the global resting-state fMRI is coupled to cortical delta-power modulation^4,5^ and pupil diameter change^37^ of a similar timescale (∼10-20 seconds in humans). It takes the form of sequential activations in the cortex mostly between the task-negative (i.e., the default mode network) and task-positive regions that often show opposite responses to task performance^7,38^.

The functional relevance of this resting-state global brain activity remains unclear, but speculations can be made with considering existing literature. First, this global activity is likely related to memory function due to its co-modulation with the hippocampal ripples. The hippocampal ripples have been found to be co-modulated, on a slow timescale of seconds, with cortical delta power^24,39^, sleep spindles^40^, cortical membrane potentials, and pupil micro-dilations^32^. There were also evidence for slow co-variation of the pupillometry and cortical neuronal activity during quiet wakefulness^41,42^. Our findings here extend these previous observations by showing that the slow modulations of the hippocampal ripples and other neural and physiological changes actually are embedded in the highly organized global activity involving the majority of neuronal population. This highly structured activity may provide local network dynamics critical for generating hippocampal ripples^39^ and also a time window for their interaction with a large population of distant neurons, particularly those in the cortex.

This resting-state global activity may also represent a specific entity enabling an off-line interaction between the memory function and cholinergic arousal system. A close relationship between the cholinergic system and memory function, particularly their concurrent impairments in the Alzheimer’s disease^43,44^, has long been evident^45,46^, and various theories have been proposed to explain this relationship^47^. The global resting-state fMRI activity has been linked to the cholinergic region at the basal forebrain^4,5^, since pharmaceutically de-activating the region greatly suppressed this global fMRI component^5^. Spontaneous low-frequency pupil diameter change during immobile rest, similar to what we observed here, has also been linked to the activity of corticopetal cholinergic neurons from the basal forebrain^42^. Together with its potential relationship with memory function as discussed above, the global resting-state brain activity may represent an “off-line” process that links the memory consolidation and cholinergic system, which is distinct from the “online” influence of the cholinergic system on memory encoding through the attention regulation^47^. This “offline” interaction is presumably mediated by extensive neuronal connections through the septohippocampal pathway linking the basal forebrain and hippocampus^48,49^.

Lastly, the resting-state global activity may orchestrate low-frequency (∼0.1 Hz) physiological modulations that are important for peri- and para-vascular drainage of brain waste. The available data only allowed us to see spontaneous low-frequency pupil micro-dilations in the present study, but other sympathetic responses may concur. The global resting-state fMRI activity has been associated with slow (∼10 sec) modulation of various physiological signals, including respiration^13,23^, cardiac pulsation^12,14,22^, heart rate variability^50^, and arterial tone^15^, presumably mediated through sympathetic innervation^14^. These low-frequency physiological modulations could be important for the clearance of toxic brain proteins by driving cerebrospinal fluid flow^51–54^ through the perivascular and interstitial spaces through a newly-discovered glymphatic pathway^17,55^. Consistent with this notion, the low-frequency spontaneous vasomotion was recently linked to paravascular clearance in mice^18^. In human, the global resting-state fMRI activity indeed was found to be coupled by cerebrospinal fluid movements^16^. Moreover, the reduction of this coupling has been found to correlate with various AD pathology, particularly increased accumulation of amyloid-beta protein^19^. Altogether, the resting-state global activity and associated physiological modulations may represent a highly coordinated neuronal and physiological process that is potentially related the cholinergic deficiency, memory dysfunction, and toxic protein accumulation, the three major pathological features of AD. It remains a challenge for future studies to further test this hypothesis and understand the role of this resting-state process in the neurodegenerative diseases and dementia.

## Acknowledgements

This work was supported by the National Institutes of Health (NIH) Pathway to Independence Award (K99/R00 5R00NS092996-03), the Brain Initiative award (1RF1MH123247-01), the NIH R01 award (1R01NS113889-01A1), and the Intramural Research Program of the National Institute of Mental Health (ZIAMH002896).

## Author contributions

**X.L**. contributed to the conception, design of the work, and data analysis;

**X.L**. also devoted the efforts to the supervision, project administration and funding acquisition;

**X.L**., **D.A.L. &Y.Y** contributed to data visualization, and writing the paper.

## Data and materials availability

All the multimodal data are available at https://alleninstitute.org. All the codes are available from the corresponding author upon request.

**Table S1.**
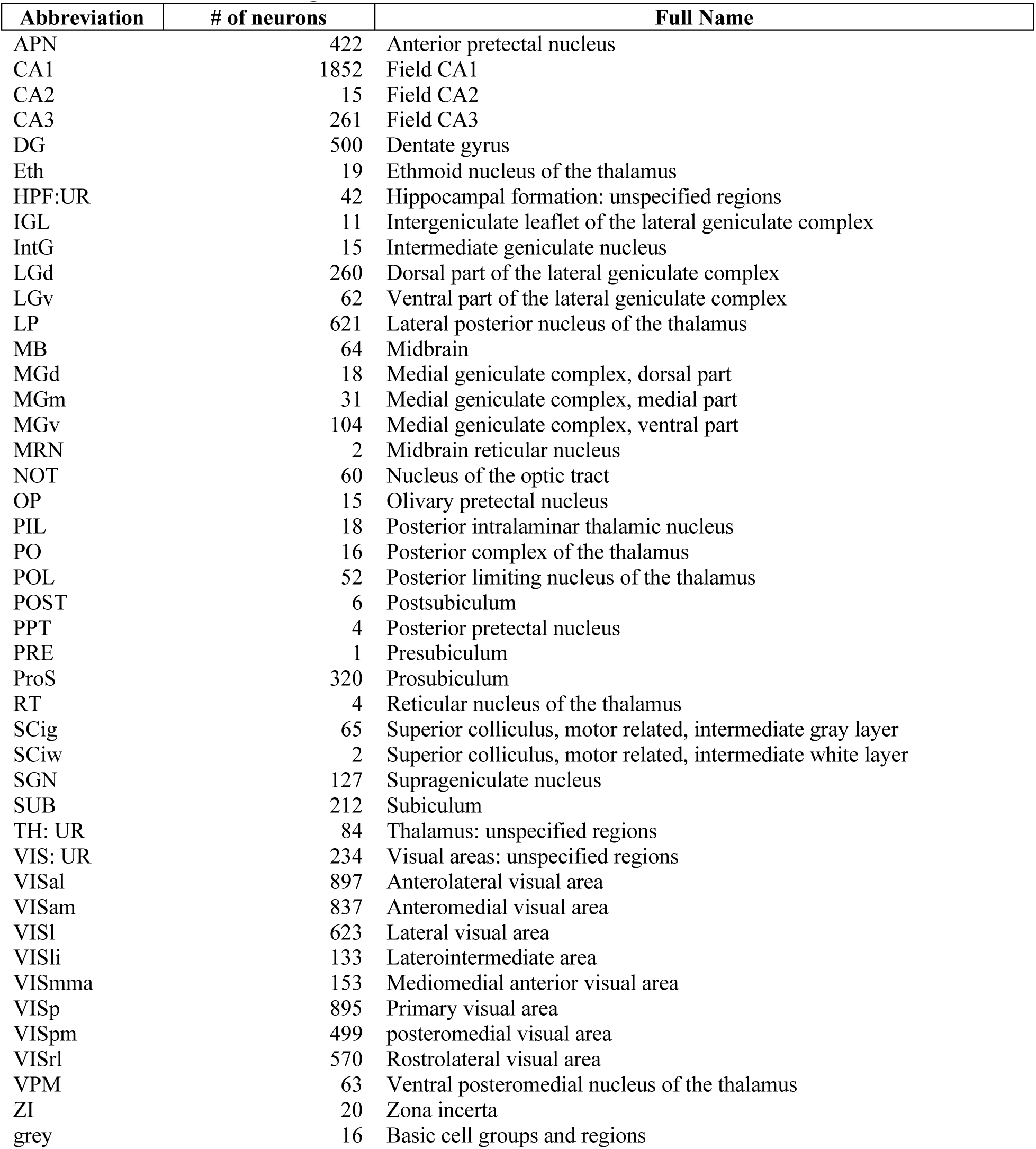
A list of brain regions with recorded neurons and their abbreviation.

**Supplementary Figure 1.**
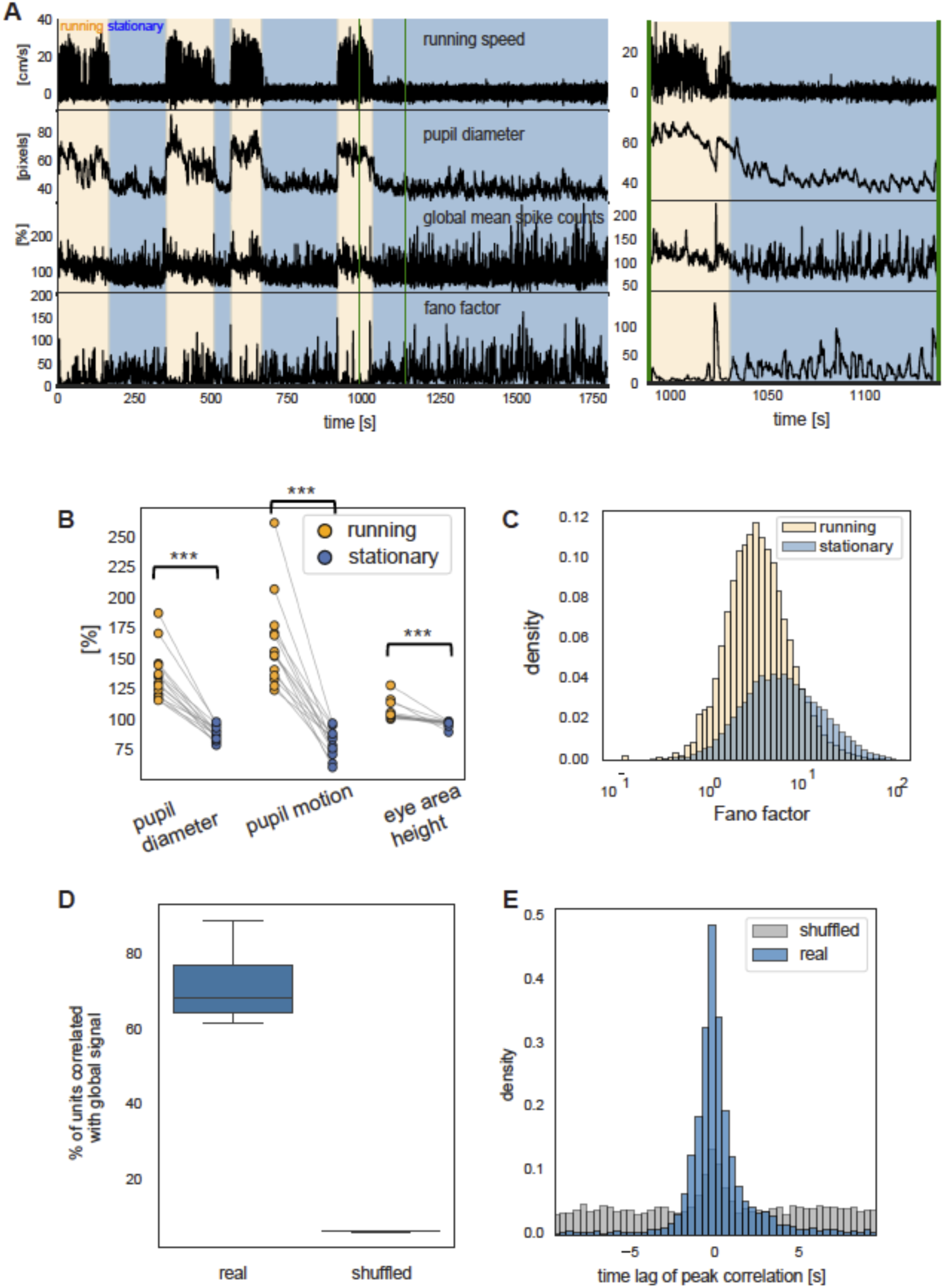
Neural and behavioral signals during the running and stationary periods. (A) Neuronal and behavioral signals (left: the entire session; right: an expanded segment) from the representative mouse showed distinct patterns during the running (yellow background) and stationary (blue) periods. The y-axis scale was re-adjusted for the expanded segment. (B) The mean pupil diameter, pupil motion, and eye-area height were summarized and compared between the running (yellow) and stationary (blue) periods in all 14 mice. (C) The Fano factor distributions for the running and stationary periods of all 14 animals. (D) The percentages of neurons whose spiking activity is significantly correlated with the global mean spiking rate across all 14 mice (blue, left). The threshold for significance (*p* < 0.05) was established based on the correlations of the phase-shuffled signals with their mean, which thus show exactly a 5% value (gray, right). (E) The distribution of the time lag where the peak correlation between individual neurons and global mean were achieved for both the real data (blue) and phase-shuffled controls (gray).

**Supplementary Figure 2.**
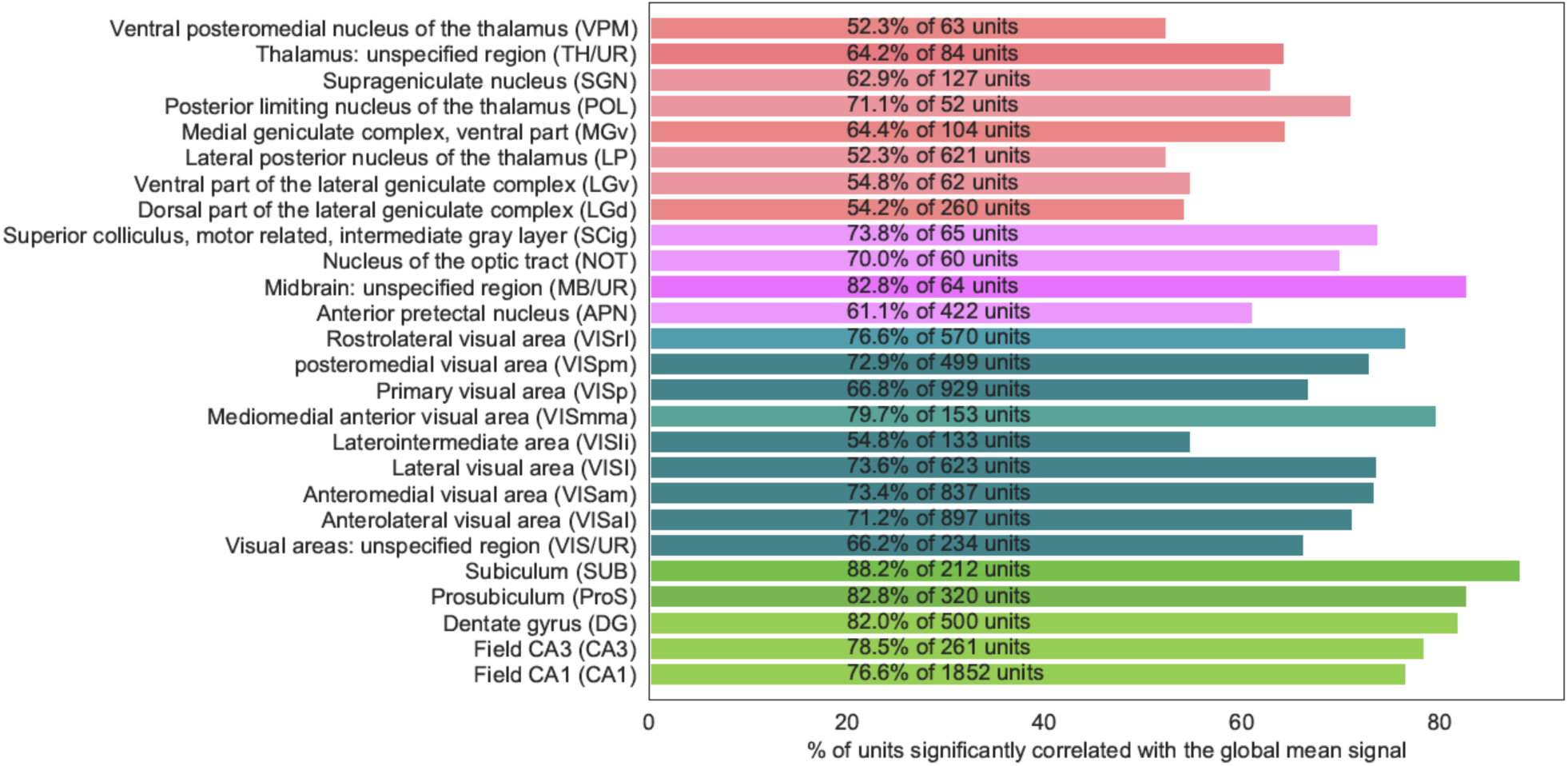
The percentages of neurons showing a significant (*p* <0.05) correlation with the global mean spiking activity in a total of 26 regions that have at least 50 neurons. The data from all the 14 mice was pooled to generate this result.

**Supplementary Figure 3.**
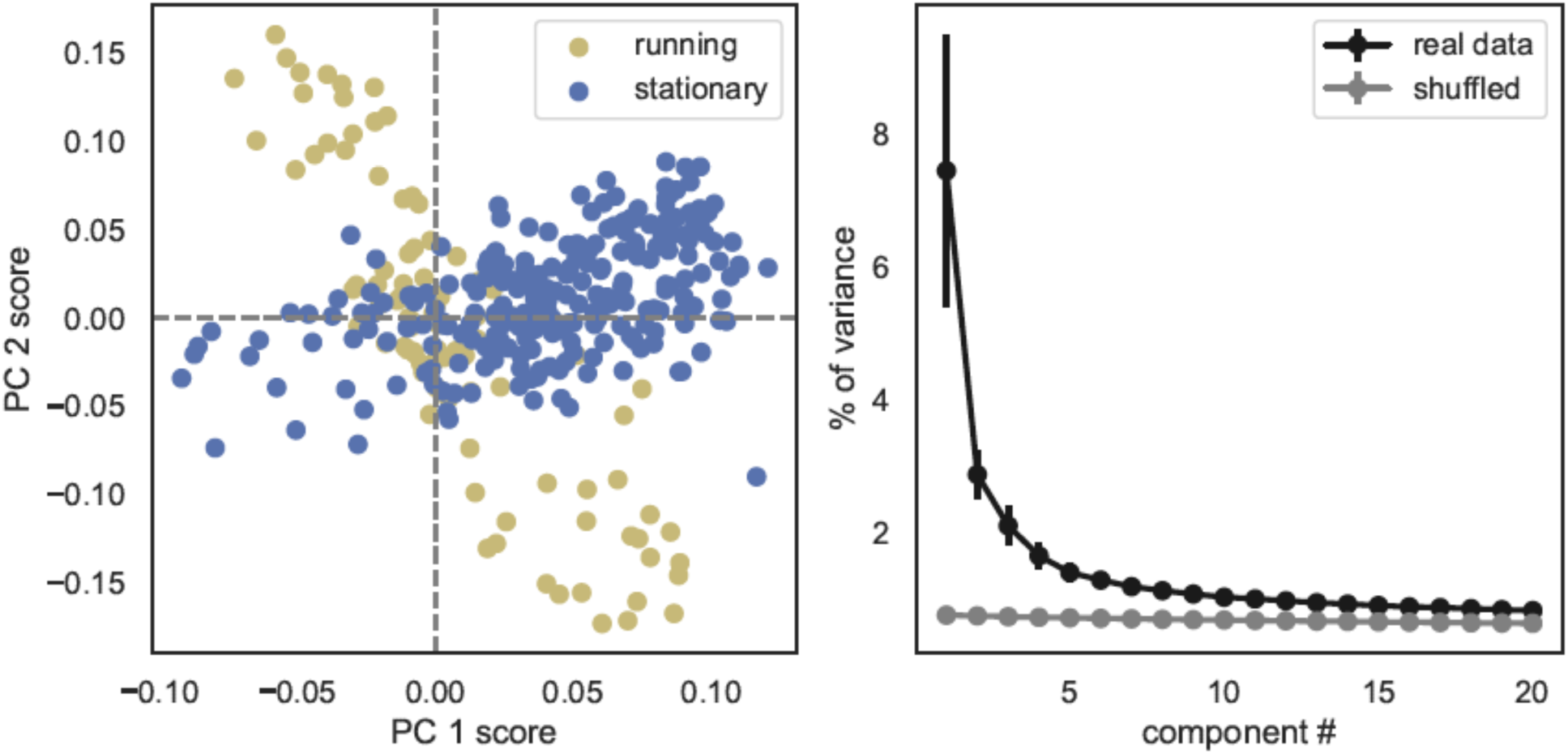
Outcomes of the delay-profile decomposition method. (A) The delay profiles of the stationary (blue) and the running (yellow) periods from the representative mouse were displayed in 2D spaces spanned by their first (i.e., the principal delay profile) and second principal components. (B) The scree plot showing the percentage of variance explained by the first 20 principal components derived from both the real data (black) and the shuffled delay profiles (gray). The error bars represent the standard deviation across the 14 mice.

**Supplementary Figure 4.**
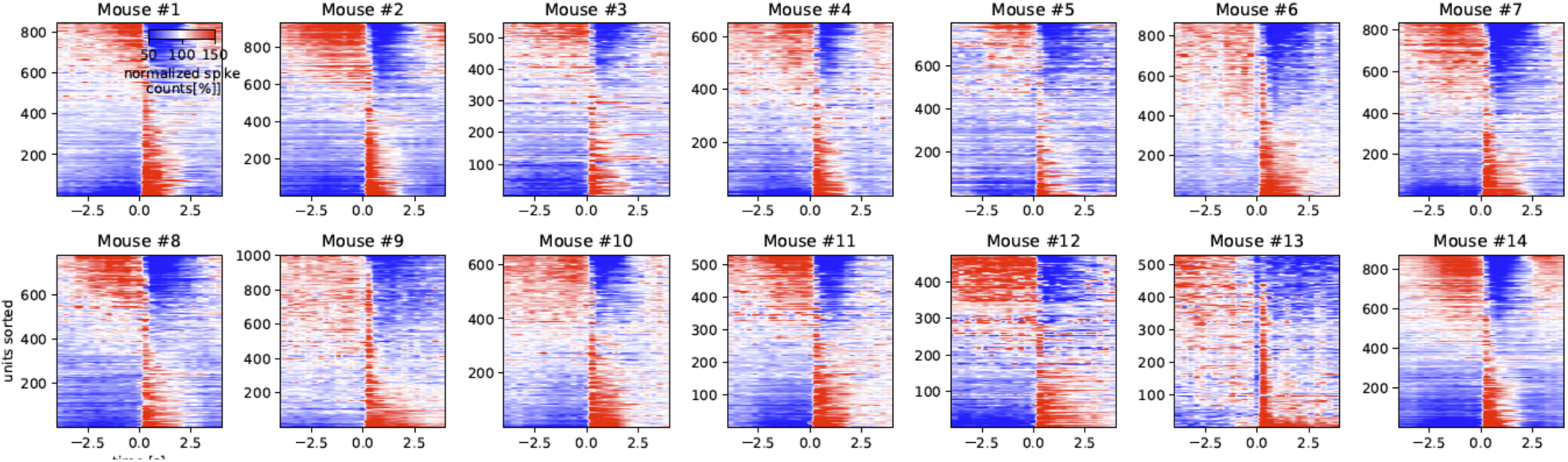
The mean patterns of sequential activations in all 14 animals.

**Supplementary Figure 5.**
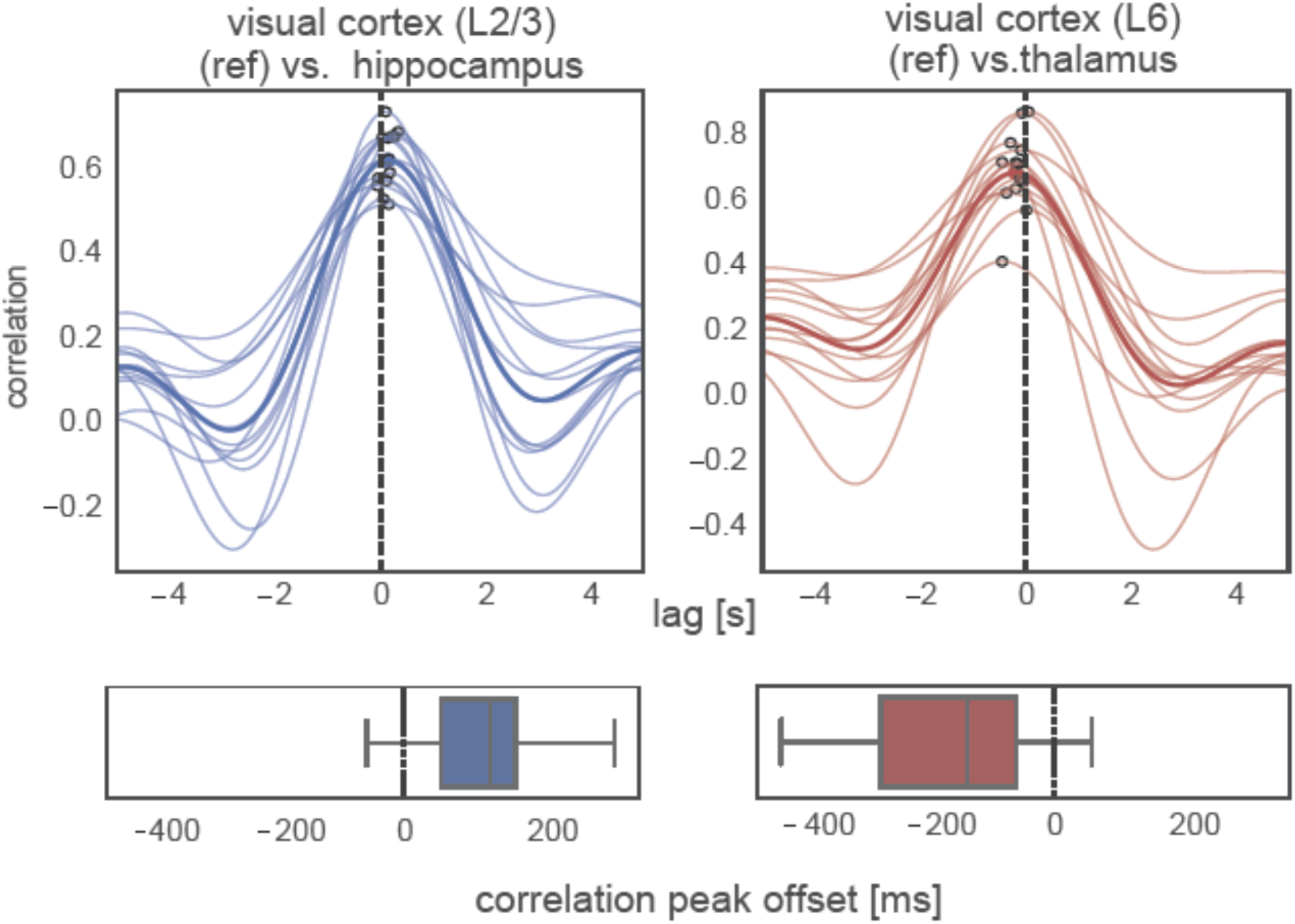
Cross-correlation functions of spontaneous spiking between the leading negative-delay neurons at the hippocampus (CA1 and DG) and the visual L2/3 layer (reference) (top left) and the distribution of peak correlation offsets (bottom left). The results were also computed between the lagging positive-delay neurons at the thalamus structures and in the L6 of the visual cortex (reference) (right). Thin and thick lines present the results of individual animals and their average, and the peak correlations are marked as hollow circles. The boxplots at the bottom display the distribution of the time lag of peak correlations.

**Supplementary Figure 6.**
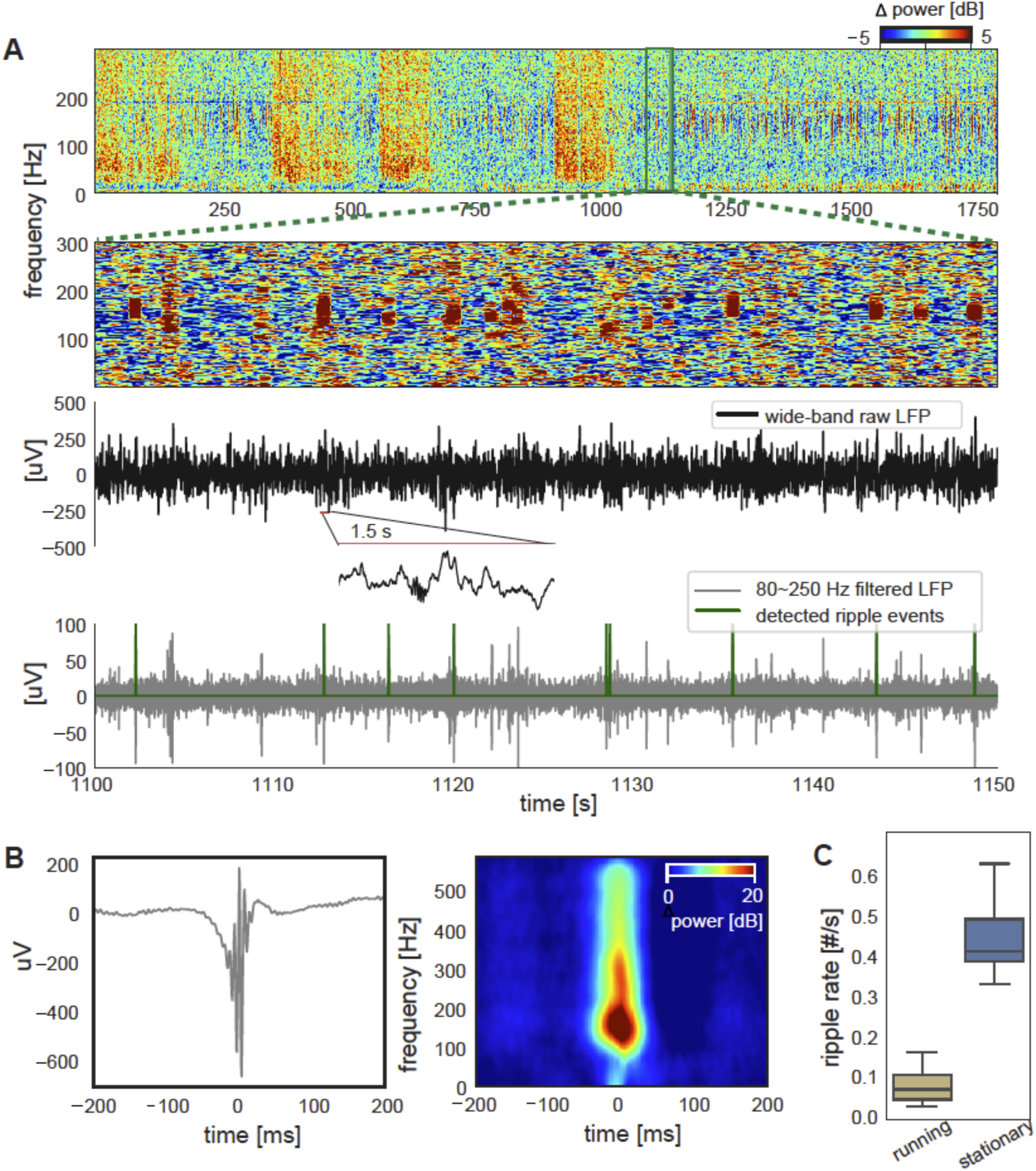
Hippocampal ripple detection. (A) Spectrogram of local field potential (LFP) signals recorded from a CA1 channel in the representative mouse shows distinct patterns in the running and stationary periods (top), with the latter being characterized by brief power bursts between 100 Hz and 200 Hz. They are clearer in an expanded 40-sec fragment of the spectrogram (the second row) and corresponding to ripple activity in the raw (the third row) and band-pass filtered (80-250 Hz) LFPs (bottom, gray), and were detected using a previously described algorithm (bottom, green). (B) The ripple activity became even clearer after being averaged across all events detected in the representative mouse, either in the time domain (left) or time-frequency domain (right). (C) The occurrence rate of the hippocampal ripples summarized for the running (yellow) and stationary (blue) periods respectively, which are significant different from each other (*p* = 3.2*×*10^−19^, paired t-test, *N* =14).

**Supplementary Figure 7.**
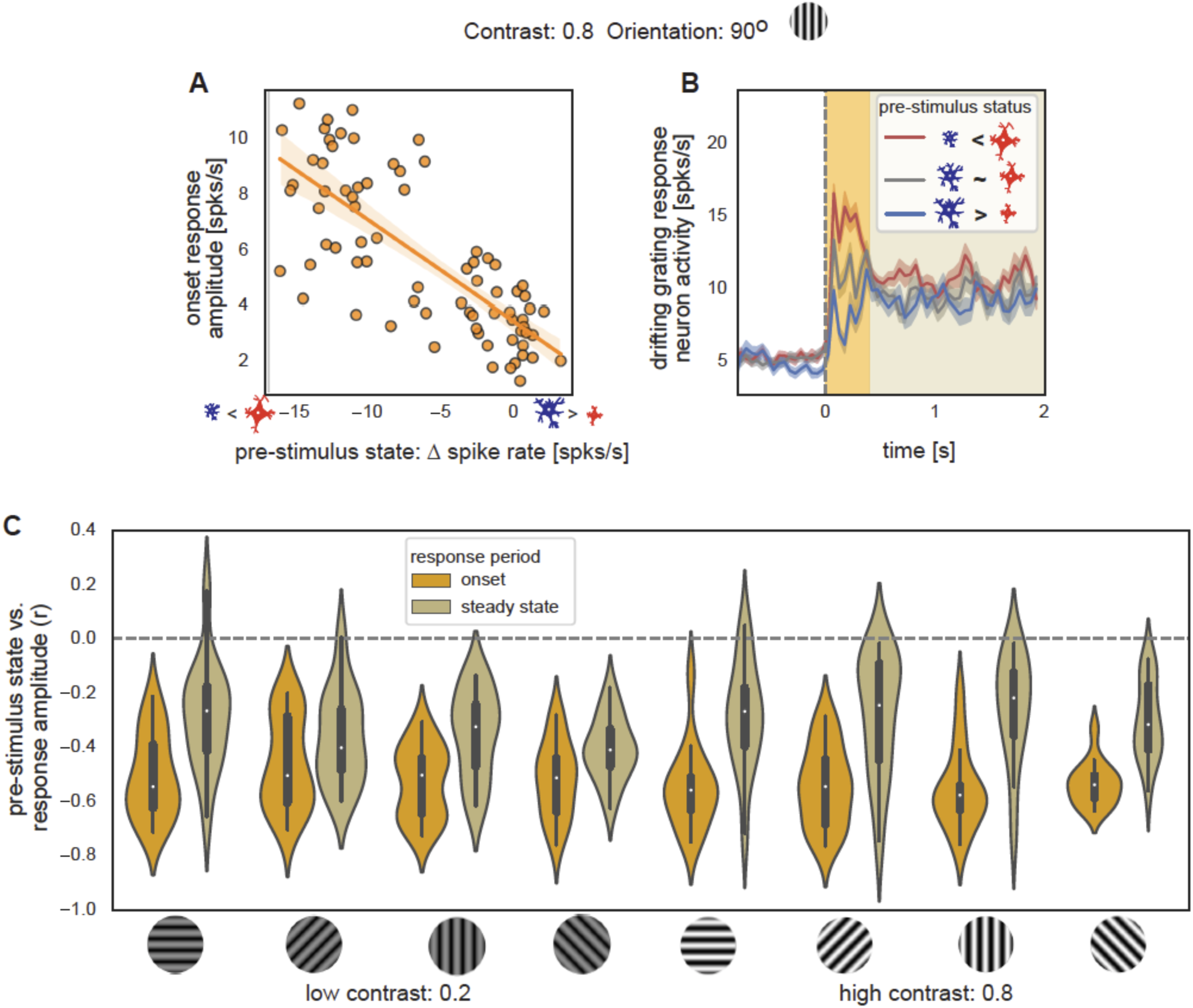
The association between the pre-stimulus state and the response amplitude. (A) The pre-stimulus state is significantly correlated (*r* = -0.76, *p* = 3.4*×*10^−15^) with the onset response amplitude to a drifting gratings stimulus (orientation: 90 degree; contrast: 0.8) across 75 trials pooled in the representative mice (left). (B) Spiking rate of the drifting gratings responding neurons were averaged within three groups of trials divided based on the pre-stimulus state: the bottom 25 trials (red) with the positive-delay neurons more active, the middle 25 trials (gray) with similar spiking rate between the negative- and positive-delay neurons, and the top 25 trials (blue) showing more active negative-delay neurons. Shadows represent regions within one SEM across trials. (C) A violin plot shows the distributions of the correlation between the pre-stimulus state and the response amplitude of the onset (orange) and steady-state (olive) stimulation periods and also for all 8 stimuli (2 contrasts *×* 4 directions).

## Materials and Methods

### Neuropixels data

We used neural and behavioral signals recorded from mice by the Visual Coding – Neuropixels project of the Allen Brain Observatory at the Allen Brain Institute^29^. The project used high-density extracellular electrophysiology probes to record spikes from a large population of neurons across a wide variety of brain areas in awake mice, including the visual cortex, hippocampal formation, and thalamus. Running and eye-tracking data were simultaneously recorded with the neural signals. The multimodal data were recorded using two sets of stimulations in two groups of mice. We focused on the “Functional Connectivity” stimulus set, which included a 30-minute “spontaneous” session without any visual stimulation. Among 26 mice of the “Functional Connectivity” stimulus set, we excluded two that had no eye-tracking data and ten with less than 10 minutes of immobile periods. The remaining 14 mice remained stationary for 22.7*±*4.3 min (range: 15.4–29.7 min) on average during the 30-minute session. Three of the 14 mice were from transgenic Cre lines (Sst-IRES-Cre, Vip-IRES-Cre, and Pvalb-IRES-Cre lines respectively), and the rest were all wild type.

We also analyzed neuronal recording data from another session with the drifting grating stimulation to examine the interaction between the ongoing and evoked spiking activities. During this session, the drifting grating stimuli were presented at a spatial frequency of 0.04 cycles/degree and a temporal frequency (2 Hz). There were eight types of drifting grating stimulus at 4 orientations (0 °, 45 °, 90 °, 135°) and 2 contrasts (10%, 80%) that were presented in random order. Each type was presented 75 times. Each trial included a 1-second baseline period without stimulation followed by a 2-second stimulation period.

Analyses of the data were conducted in Python with the use of Allen Software Development Kit (SDK) (https://allensdk.readthedocs.io/en/latest/).

### Definition of the running and stationary states

The running and stationary states were defined based on the running speed. The running speed signal was first temporally filtered with a low-pass filter (3rd-order Butterworth filter with critical frequency at 0.05 Hz) to reduce the high-frequency noise. Negative values of the running speed data resulted only from noise. We identified 99.95 percentile of all negative values, and took its absolute value as a threshold, and classified a time point with a running speed higher than this value as the running period time point. We used different thresholds, ranging from 95 to 99.99 percentile, and the classification results are largely similar and consistent with the visual perception based on the running speed signal. The resulting running period was often intermixed with short and brief stationary epochs. We thus re-classified the stationary periods shorter than 50 seconds to the running period to have the stationary periods represent a more extended period of the immobile state. As a result, the running state includes a small portion of short immobile epochs.

### Eye data

The eye-tracking data were pre-processed by the Neuropixels project and a set of parameters were computed based on images recorded by eye-tracking camera at a sampling rate of 30 Hz. The procedure was described in detail in the white paper of the Neuropixels project. We computed multiple eye/pupil indexes based on these parameters. The pupil diameter was defined as the mean of the pupil height and width. The eye motion was defined as the distance by which the pupil center moved between two consecutive time points. We also used the height of the eye area summarized by the Neuropixel project.

### Neuronal spikes

We directly used neural spikes sorted by the Neuropixels project. Kilosort2^28^ was used for spike-sorting and details were included in the white paper of the project. Spikes of each neuron were counted every 200ms to obtain a time course for spiking activity. The spiking time course of individual neurons was normalized by its temporal mean and then averaged to obtain the global mean spiking activity. The normalization was conducted to equalize the contribution from fast-spiking and slow-spiking neurons. A second version of the global mean spiking signal was also computed without this normalization procedure, and the two versions were however highly similar to each other.

Cross-correlation functions were computed between the spiking activity of individual neurons and the global mean spiking activity. The spiking signals were first filtered by a lowpass filter (<0.5 Hz) to focus on low-frequency components. The correlations were calculated at time lags between *±*10 seconds. The peak (of the absolute value) and corresponding lag of each cross-correlation function were identified and recorded. We then repeated the identical procedure after shuffling phases but preserving the frequency profile of each neuron’s spiking time course. In other words, we correlated the phase-shuffled signals to their own mean and recorded the peak absolute correlation and corresponding lag. The phase-shuffled frequency-profile-preserved controls were used to account for the portion of correlations that may arise from the low-frequency filtering and averaging. The peak absolute correlations between the phase-shuffled signals and their mean were then regarded as a null distribution, and its 95% percentile was used as a threshold to determine whether individual neurons’ spiking activity is significantly (*p* < 0.05) correlated with their global mean. The number of neurons showing significant correlations with the global mean spiking activity was counted and summarized for all 14 animals. Such data were also pooled across animals and summarized separately for 26 brain regions with more than 50 recorded neurons across all the mice.

### Fano Factor

The Fano factor was computed to quantify the high-order correlation and global synchronization level of the population spiking activity^30,31^. The Fano factor was defined as the ratio between the variance and the mean of population spike count, i.e., the sum of spikes across the neuronal population within each 200 ms time bin.

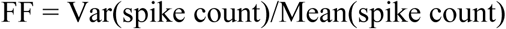

The variance and mean were computed within a 2-sec time window that was slid over time by a step of 200ms, generating a continuous time course for the Fano factor. Synchronized population events would lead to a temporary surge of population activity, which in turn would increase the variance of population activity and thus the Fano factor. We summarized and compared the distribution of the Fano factor values between the running and stationary periods. The effect of the sliding window size on the Fano factor was also estimated by using different window sizes ranging from 1 second to 6 seconds with a 1-second increment. The Fano factor computed with a large sliding window appeared to be a temporally lowpass filtered version of that obtained by a small sliding window. The window size would however not affect the overall pattern of the Fano factor nor its significant difference between the running and stationary states.

### Principal delay profile

We adapted a delay profile decomposition method^7^ to derive the principal delay profile of the neuronal spiking activity, which should map the major direction of sequential activations across the neuronal population if any exists. This method has been successfully used to extract the propagating activity of rsfMRI signals and also validated with simulation^7^. Briefly, we cut the neuronal spiking data into time segments based on troughs of the lowpass filtered (<0.5 Hz) global mean spiking signal, so each segment contained a single peak of the global mean spiking signal. For each time segment *j*, we constructed a delay profile *d*_*j*_whose entry *d*_*“j*_is the timing of the spiking count centroid of neuron *i* within this time segment. Each delay profiled was then centered and thus represented the relative delay of each neuron’s spiking activity with respect to the overall population mean with this specific time segment. The use of centroid timing can be considered as an improvement to the original approach that used the local peak timing by avoiding the filtering procedure on original signals. Overall, this new approach generated similar but more robust results as compared with its original version. The delay profiles of all M segments were then combined into a delay profile matrix D whose the j-th column is the delay profile for the time segment *j*. We standardized delay matrix columnwise and then applied singular value decomposition (SVD) to derive the principal delay profile:

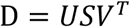

The principal delay profile *d*^*∗*^ was defined as the first column of U, the singular vector corresponding to the largest singular value.

We correlated the principal delay profile *d*^*∗*^ with each delay vector *d*_*i*_, and the resulting vector of correlations is related to the first column of *V* by a scaling factor. However, this correlation vector allowed us to have a more initiative assessment of similarities between the principal delay profile and individual delay profiles. This vector of correlations was transformed to Z scores using a Fisher’s transformation. Based on it, we then identified time segments with a delay profile positively (*p* < 0.001) correlated with the principal delay profile. These time segments were regarded as showing the sequential activation. We next defined the negative- and positive-delay neurons as those whose mean delay value, averaged across the stationary time segments of sequential activation, significantly (*p* < 0.001) lower or higher than zero respectively. We also randomly shuffled the values of each delay profile and correlated the shuffle delay profiles with the principal delay profile.

The neural and behavioral signals were averaged across the time segments of sequential activation during the stationary period. These time segments were aligned according to the time point when the positive-delay neurons showed the sharpest increase in their mean spiking activity, i.e., the maximum of the 1^st^ order derivative.

The principal delay profile was rescaled to the seconds unit based on the average pattern of sequential spiking activity. The original principal delay profile from SVD is unitless. To rescale it to values in the unit of seconds, we first identified the time point when each neuron reached its peak spiking activity in the average pattern of sequential activation. We then fitted a linear regression model between the original principal delay profile and the peak timing, and the regression coefficient between the two was regarded as a scaling factor. We multiplied the original principal delay profile by this scaling factor to obtain the version in the unit of second.

We further quantified the relationship between neurons’ principal delay value and their modulation across the running and stationary state. The spiking rate difference between the running and stationary states was computed for each neuron, which was correlated with the principal delay values across all the neurons recorded in the same mouse. The correlation was further summarized across all the mice.

The neuronal population dynamics were quantified at the transitions between the running and stationary state. We located all the transitioning time points between the running and stationary states and focused only on isolated ones by excluding those with another transitioning point with a *±*40 sec time window. The transitioning points were divided into the running-to-stationary transitions and stationary-to-running transitions. The spiking data of all neurons were then aligned and averaged according to these two types of transitions separately. The mean spiking activity of negative- and positive-delay neurons, as well as the running speed, were also aligned and averaged according to the transitions in the same way.

### Delay map

The principal delays of all the neurons were pooled and mapped across all 14 mice. As described above, the values of the principal delay profile were re-scaled into the unit of seconds. The principal delays of all surveyed neurons were then pooled across all 14 mice and displayed using colors at their recording sites. For neurons recorded by the same channel, their mean delay was computed for mapping. There was a total of 6171 channels on 79 probes implanted in all 14 animals. They spanned across a relatively narrow range along the anterior-posterior direction. We thus collapsed them along this direction and mapped them onto a 2D brain image in the middle of the recording sites.

The region-specific distributions of the principal delay were also summarized. This was done for the 24 brain regions with more than 50 recorded neurons across all the animals. For the principal delays of each region, we conducted a one-sample t-test to test whether their mean was significantly different from zero, and the results were further corrected for multiple comparisons using the Bonferroni method. For regions of the visual cortex, the neurons were further subgrouped according to the cortical layers they located and the principal delays were summarized accordingly. Very few neurons were recorded from layer I, and they were thus removed from the analysis. For each layer-specific summary, the 95% conference interval for the mean principal delay was computed.

The percentages of the negative- and positive-delay neurons were summarized for the same 24 brain regions with more than 50 surveyed neurons. The identification of negative- and positive-delay neurons was described above. The region-specific ratio of these two types of neurons was computed in each brain region and compared with the overall ratio using a proportion z-test with the Bonferroni correction.

The relative time delay of spontaneous spiking activity was estimated between different brain regions among the same type of neurons, i.e., the negative- or positive-delay neurons. Specifically, we examined the delay between the negative-delay neurons from the L2/3 of the visual cortex and the hippocampus (only the CA1 and DG), as well as between the positive-delay neurons from the L6 of the visual cortex and the thalamus, based on the time offset of the peak cross-correlations. For each mouse, the spiking activity of the negative neurons was averaged separately within the L2/3 of the visual cortex and the hippocampus (only the CA1 and DG regions) during the stationary state. Their cross-correlation function was computed, and the peak correlation and corresponding time lag were then identified. A similar analysis was conducted between the positive-delay neurons recorded in the visual L6 and the thalamus. The time offset of the peak correlation was then summarized across all 14 mice except one without any neurons recorded in the L2/3 of the visual cortex and the thalamus (session ID = 829720705). A one-sample t-test was used to test whether the mean time offset of the peak correlation was significantly different from zero.

### Cortical delta powers

The delta-band power was computed for local field potentials (LFP) recorded at the cortex, which was exclusively the visual cortex for this study. A band-pass filter (0.5-4 Hz) was first applied to the wide-band LFP signal of each cortical channel. The filtered signal was then rectified and lowpass filtered (<0.72 Hz, which was corresponding to π cycles of the mean band-pass frequencies) to extract the delta-band power. The delta-band power was highly correlated across the cortical channels during the stationary period, and we thus took their mean to represent the cortical delta-band power dynamics.

The relative time delay between the neural and behavioral signals was estimated through their cross-correlation functions during the stationary state. Specifically, the cross-correlation functions were computed between the spiking activity of the negative-delay neurons and the cortical delta powers, as well as between the spiking activity of the positive-delay neurons and the pupil diameter, during the stationary state. The offset time of peak correlation was identified and summarized for all the animals. We then tested whether the mean of this peak correlation offset was significantly different from zero using the one-sample t-test.

### Hippocampus ripples detection

We adapted an existing algorithm of offline ripple detection^33^ to detect hippocampal ripples in LFP recorded from the CA1 region. The LFP was originally acquired at a sampling rate of 2.5k Hz but downsampled to 12.5 Hz by the Neuropixels project. Briefly, the LFP from each CA1 channel was bandpass (80-250 Hz) filtered and then clipped outside the mean *±* 5SD range to minimize the ripple-rate induced biasing. The filtered and clipped signal was then rectified and lowpass filtered (<52.5 Hz, which was corresponding to π cycles of the mean band-pass frequencies) to extract the signal power. The mean and SD were calculated for the power of the clipped signal. Then, the power of the unclipped signal was also computed, and all events exceeding 5 SD from the mean were identified as ripple events. Short events with a duration of < 15 ms were discarded, and adjacent events with a short gap of <15 ms were merged. The detected events were then expanded until the power fell below 2 SD. We aligned the ripple events according to the trough closest to the peak power and then averaged them directly in the time domain or their spectrogram instead. The spectrograms were computed with a time window of 960 seconds and normalized with respect to the first 100ms baseline period (subtract the baseline power level of this period). The ripples detected in all CA1 channels were largely overlapped with each other. We defined a union mask of ripple events in which that more than 40% CA1 channels showing ripples, and identified a single channel with the highest ripple power within this mask as the representative channel.

### Drifting grating responses

We investigated how the ongoing brain dynamics affect neuronal responses to external sensory stimulation using spiking data from another 30-minute session of drifting grating stimulation. In addition to the negative- and positive-delay neurons identified above, we identified a third group of neurons that responded to the drifting gratings. The drifting-grating-responding neurons were defined as those showing a significantly (*p <* 0.001, paired t-test, *N* = 75 trials for each stimulus in each mouse) higher spiking rate during the simulation period (0-2 sec with respect to the stimulus onset) than the baseline period (–0.8-0 sec, the first 200 ms was excluded to avoid any residual effects of the previous trial). There were limited overlaps between the three groups of neurons. Nevertheless, to rule out any potential effects due to overlapped neurons, we excluded, from the negative- and positive-delay neuronal groups, any neurons showing significant (*p* <0.05; more liberal than the standard for defining the drifting grating responding neurons) changes in response to drift-grating stimulation. Likewise, we also excluded, from the drift-grating responding group, neurons whose delay profile values for the time segments of sequential activation are significantly different (*p* <0.05; more liberal than the standard by which we defined the negative- and positive-delay neurons) from zero.

We quantified the pre-stimulus state of the negative- and positive-delay neurons and the response amplitude of the drifting-grating-responding neurons. The pre-stimulus state was quantified by the spiking rate difference between the negative- and positive-delay neurons within a 200-ms time window before the stimulus onset. The response amplitude of the drifting-grating-responding neurons to drifting grating stimuli was quantified by their spiking rate difference between the stimulation and baseline periods. The response amplitude was quantified separately for an early onset phase (0-0.4 sec) and a late steady-state phase (0.4-2 sec) of the simulation period. For each of eight types of drifting grating stimuli, the pre-stimulus state was correlated with the response amplitude either across 75 trials within the same animal or 1050 trials pooled across all 14 animals. We also characterized the spiking dynamics of the drifting-grating-responding neurons over trials with distinct levels of pre-stimulus state. For each mouse and each drifting grating type, we divided 75 trials into three groups according to tertiles of the pre-stimulus state values, and simply averaged the spiking activity of the drifting-grating-responding neurons within each group. The spiking dynamics was further averaged across all animals for the same type of drifting grating stimuli.

